# Bacterial amino acid auxotrophies enable energetically costlier proteomes

**DOI:** 10.1101/2025.01.24.634666

**Authors:** Niko Kasalo, Tomislav Domazet-Lošo, Mirjana Domazet-Lošo

## Abstract

The outsourcing of amino acid (AA) production to the environment is relatively common across the tree of life. We recently showed that the massive loss of AA synthesis capabilities in animals is governed by selective pressure linked to the energy costs of AA production. Paradoxically, these AA auxotrophies facilitated the evolution of costlier proteomes in animals by enabling the increased use of energetically expensive AAs. Experiments in bacteria have shown that AA auxotrophies can provide a fitness advantage in competition with prototrophic strains. However, it remains unclear whether energy-related selection also drives the evolution of bacterial AA auxotrophies and whether this affects the usage of expensive AAs in bacterial proteomes. To investigate these questions, we computationally determined AA auxotrophy odds across 980 bacterial genomes representing diverse taxa and calculated the energy costs of all their proteins. Here, we show that auxotrophic AAs are generally more expensive to synthesize than prototrophic AAs in bacteria. Moreover, we found that the cost of auxotrophic AAs significantly correlates with the cost of their respective proteomes. Interestingly, out of all considered taxa, Mollicutes and Borreliaceae—chronic pathogens highly successful in immune evasion—have the most AA auxotrophies and code for the most expensive proteomes. These findings indicate that AA auxotrophies in bacteria, similar to those in animals, are shaped by selective pressures related to energy management. Our study reveals that bacterial AA auxotrophies act as costly outsourced functions, enabling bacteria to explore protein sequence space more freely. It remains to be investigated whether this relaxed use of expensive AAs also enabled auxotrophic bacteria to evolve proteins with improved or novel functionality.

## 1. Introduction

The concept of functional outsourcing suggests that organisms streamline their genomes by losing costly functions and substituting them through biological interactions [1]. A particularly striking example is the outsourcing of amino acid (AA) production in animals, which almost universally lack the ability to synthesize about half of the proteinogenic AAs [1–6]. Our recent research demonstrates that this phenotype is driven by selective pressures related to the energy cost of AA production, allowing animals to evolve proteomes that incorporate expensive AAs more frequently [2]. These findings raise the possibility that similar correlations between proteome costs and AA biosynthetic capabilities exist in other major clades on the tree of life, including bacteria.

Compared to animals, the understanding of bacterial AA biosynthesis remains relatively incomplete. Earlier studies indicated that most bacteria possess at least a few auxotrophies [7,8]. However, recent research has revealed that the prevalence of AA auxotrophies may have been overestimated due to gaps in knowledge regarding bacterial biosynthetic pathways, though many bacteria still lack the ability to synthesize the full set of AAs [4,9]. Experimental evidence shows that AA auxotrophies can confer a fitness advantage in competition with prototrophic strains [7,10], and they may serve as an evolutionary strategy to reduce biosynthetic burdens via cooperative interactions within microbial communities [11].

However, the adaptive benefits of AA auxotrophies in bacteria remain debated. A previous study speculated that outsourcing AA biosynthesis might reduce cellular metabolic costs [10]. While it is well-established that AAs vary in biosynthetic cost [11–13] and that costlier AAs promote stronger microbial cross-feeding interactions [11], a recent study found no significant correlation between AA biosynthesis costs and the prevalence of AA auxotrophies among bacteria [4].

Nevertheless, alternative approaches could provide a more rigorous test of whether energy-related selection drives the evolution of AA auxotrophies. For instance, if such selection influences the loss of AA biosynthetic capabilities, it should also be reflected in the incorporation of more expensive AAs into bacterial proteomes [2]. To our knowledge, no study has yet examined bacterial proteome costs in this context.

Detecting auxotrophies remains an active area of research, employing various methodologies. Many studies rely on in silico approaches, such as genome-scale metabolic modeling [14,15] or homology-guided enzyme annotation [4,7,16]. To assess their accuracy, these computational methods are typically validated against a limited number of experimentally derived datasets [4,14]. However, no simple or standardized protocol currently exists for comparing AA auxotrophy estimates across studies.

To address these gaps, we assembled a taxonomically diverse dataset of bacterial proteomes predicted from genome sequences and examined whether AA biosynthesis costs explain trends in AA auxotrophy composition. Using a simple auxotrophy detection methodology based on the MMseqs2 clustering pipeline, we obtained results comparable in quality to previous approaches. Our findings reveal that costlier AAs are more frequently lost and that bacteria with more expensive AA auxotrophies encode costlier proteomes, mirroring patterns observed in animals. These results suggest that energy-driven selection plays a key role in shaping AA auxotrophies in bacteria, enabling them to explore protein sequence space more freely during evolution.

## 2. Results

### 2.1. Global trends in bacterial AA auxotrophies

To explore global patterns in the evolution of bacterial auxotrophies, we assembled a database of 980 high-quality proteomes, computationally inferred from their corresponding genomes, capturing broad bacterial diversity (Table S1). We assessed the completeness of AA biosynthesis pathways in these species using an MMseq2-based clustering approach [17]. Our method clusters, in a single step, a representative sample of bacterial enzymes known to catalyze reactions in AA biosynthesis pathways together with all bacterial proteomes in our database (see Materials and Methods). Based on the composition of the recovered clusters, functional information on AA anabolism is then transferred between cluster members, allowing us to determine the completeness of 20 AA biosynthesis pathways for each species. Unlike previous studies that rely on strictly defined cutoff values to designate auxotrophies—thereby losing part of the available information—we analyzed pathway completeness values directly. This approach provides a more accurate representation of the likelihood that a particular pathway is present.

For each of the twenty amino acids, completeness scores range from 0 to 1, where 0 indicates that all enzymes in the pathway are absent, and 1 indicates that all enzymes are present in a given species (Fig. 1, File S1). When summing the completeness scores for all 20 AAs, we found that the average value across all bacterial species is 17.33, and that nearly 66% of species in our dataset exhibit total pathway completeness above 19. These high completeness values suggest that many bacteria are prototrophic for most amino acids, aligning with findings from recent studies [4,9]. However, certain taxonomic groups—including Lactobacillales, Mollicutes, and Borreliaceae—exhibit substantial incompleteness in multiple amino acid biosynthetic pathways (Fig. 1, File S1), indicating that auxotrophies are common in these groups.

**Figure 1.**
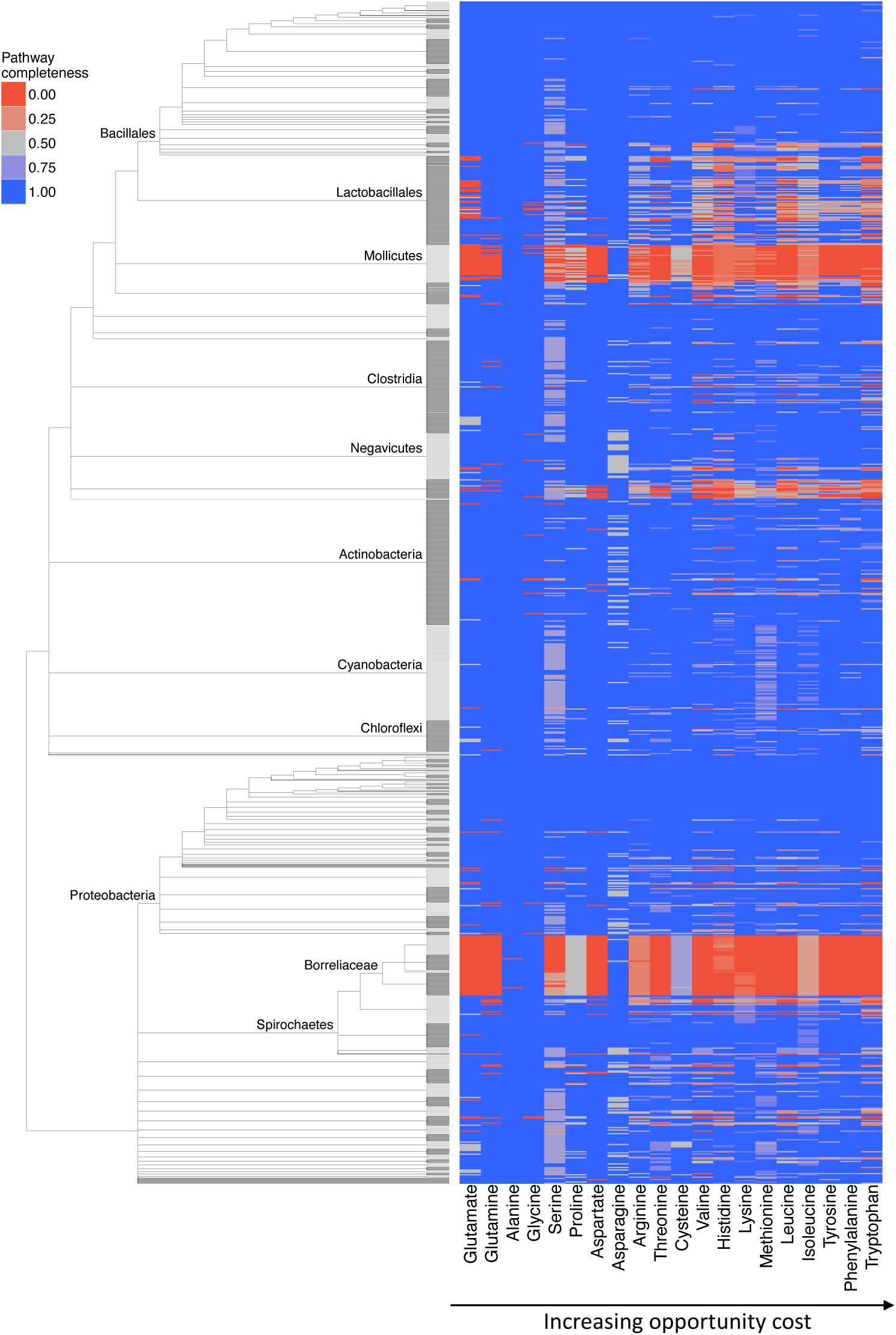
Completeness of AA biosynthesis pathways in bacteria. We created a database of 980 bacterial species to get a comprehensive overview of AA dispensability in this group. Fully resolved tree is shown File S1. We retrieved data on enzymes involved in AA biosynthesis pathways from the KEGG and MetaCyc databases and searched for their homologs within our reference database using MMseqs2 clustering (see Methods). For each AA, we showed a completeness score (CS), which represents the percentage of enzymes within a pathway that returned significant sequence similarity matches to our reference collection of AA biosynthesis enzymes. In the case of AAs with multiple alternative pathways, we showed the results only for the most complete one.

### 2.2. Expensive AAs are more commonly lost

If energy-driven selection influences the evolution of bacterial AA auxotrophies, one would expect the ability to synthesize energy-expensive AAs to be lost more frequently. To test this hypothesis, we devised an AA auxotrophy index (AI, Equation 1), defined as 1 minus the completeness score, and compared it against the opportunity cost [2], which estimates the energy expenditure associated with AA synthesis (see Materials and Methods). We observed a significant moderate correlation between opportunity cost and auxotrophy index when opportunity cost values for respiratory metabolism were applied (Fig. 2A, B). However, under fermentative conditions, this correlation was much weaker and not statistically significant (Fig. 2C). Together, these findings suggest that the evolution of AA auxotrophies in bacteria, similar to that in animals, is primarily driven by selection favoring energy savings during AA synthesis—a pattern most pronounced under respiratory conditions.

**Figure 2.**
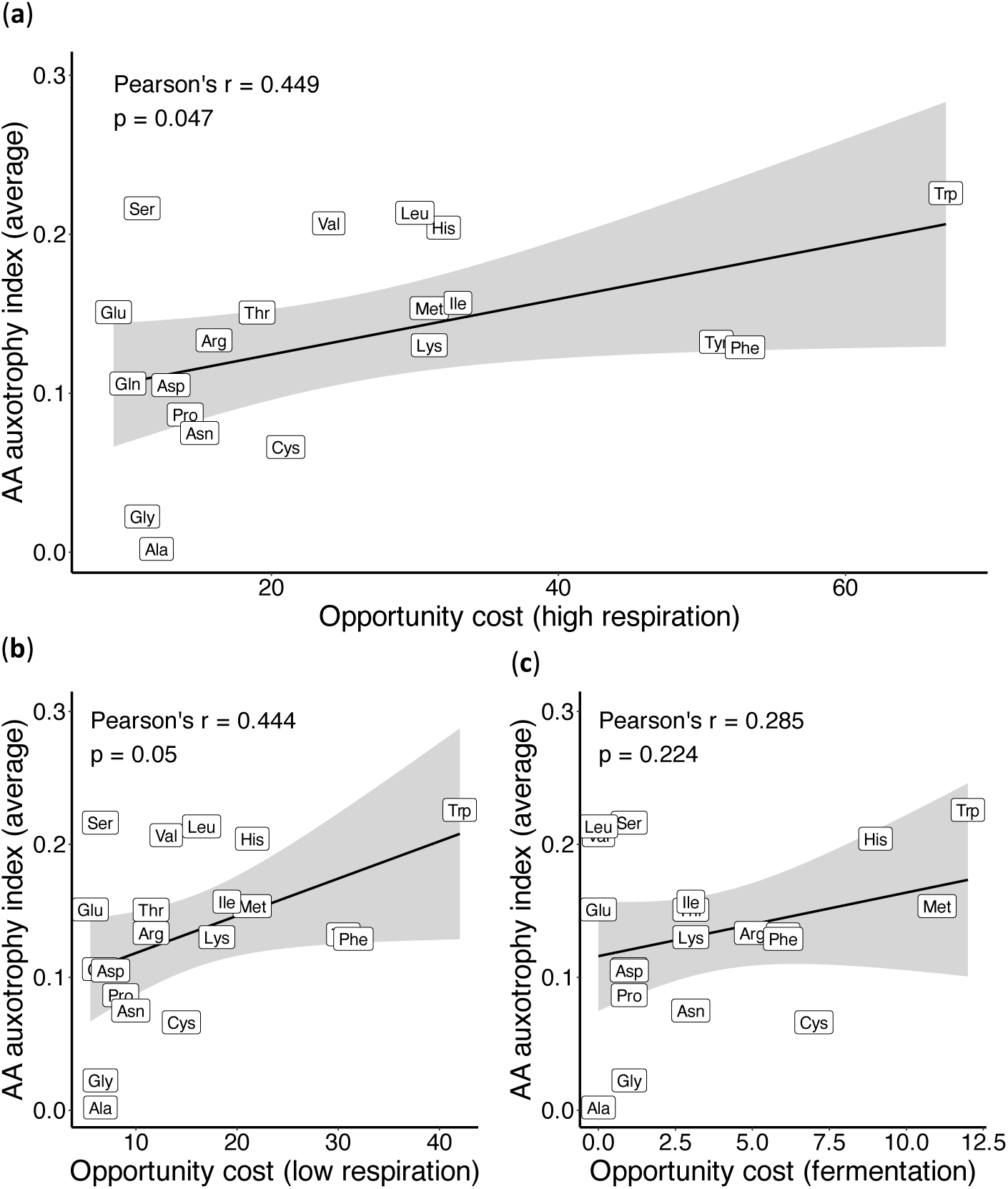
Correlation between AA biosynthesis cost and the average AA auxotrophy index. We estimated the AA auxotrophy index (AI), defined as 1 minus the completeness score (CS), for 980 bacterial species and calculated the average AI value for each AA. This value was then correlated with the opportunity cost of each AA, previously calculated for three different respiratory modes: high respiration, low respiration, and fermentation [2] (see Materials and Methods). The Pearson correlation coefficient and p-value are displayed on the graph.

### 2.3. Energy savings via AA auxotrophies enable costlier proteomes

However, the central question remains: how do reductions in AA biosynthetic pathways influence the overall cost of encoded proteomes? If energy-driven selection underpins the evolution of AA auxotrophies in bacteria, one would expect auxotrophic species to maintain more expensive proteomes than their prototrophic counterparts. This is because AA auxotrophs expend considerably less energy on AA biosynthesis and acquire the missing AAs at a relatively low cost from the environment. In essence, this pattern suggests that the energy saved on synthesizing costly AAs offsets part of the proteome’s energy expenditure, enabling auxotrophs to incorporate more expensive proteins—potentially contributing to novel functions [2].

To test this idea, we calculated the opportunity cost of each bacterial proteome (OC_*proteome*_, Equation 2) as well as the overall biosynthetic savings achieved through AA auxotrophy (OC_*savings*_, Equation 3). These calculations were performed across three respiratory modes: fermentation, low respiration, and high respiration [2]. We then assessed the correlation between OC_*proteome*_ and OC_*saving*_ (Fig. 3). Regardless of the respiratory mode used to estimate energy expenditure, our results showed that species achieving the greatest energy savings in AA production also maintain the most expensive proteomes. Similar to animals, bacterial auxotrophies appear to influence not only the immediate energy budget related to AA production but also reduce selective pressure against the use of energetically costly AAs in proteomes. Notably, we found that Mollicutes and Borreliaceae are almost invariably highly auxotrophic and possess extremely expensive proteomes. On the other hand, Lactobacillales exhibit a wide range of auxotrophy levels and have proteomes that are slightly more expensive than average.

**Figure 3.**
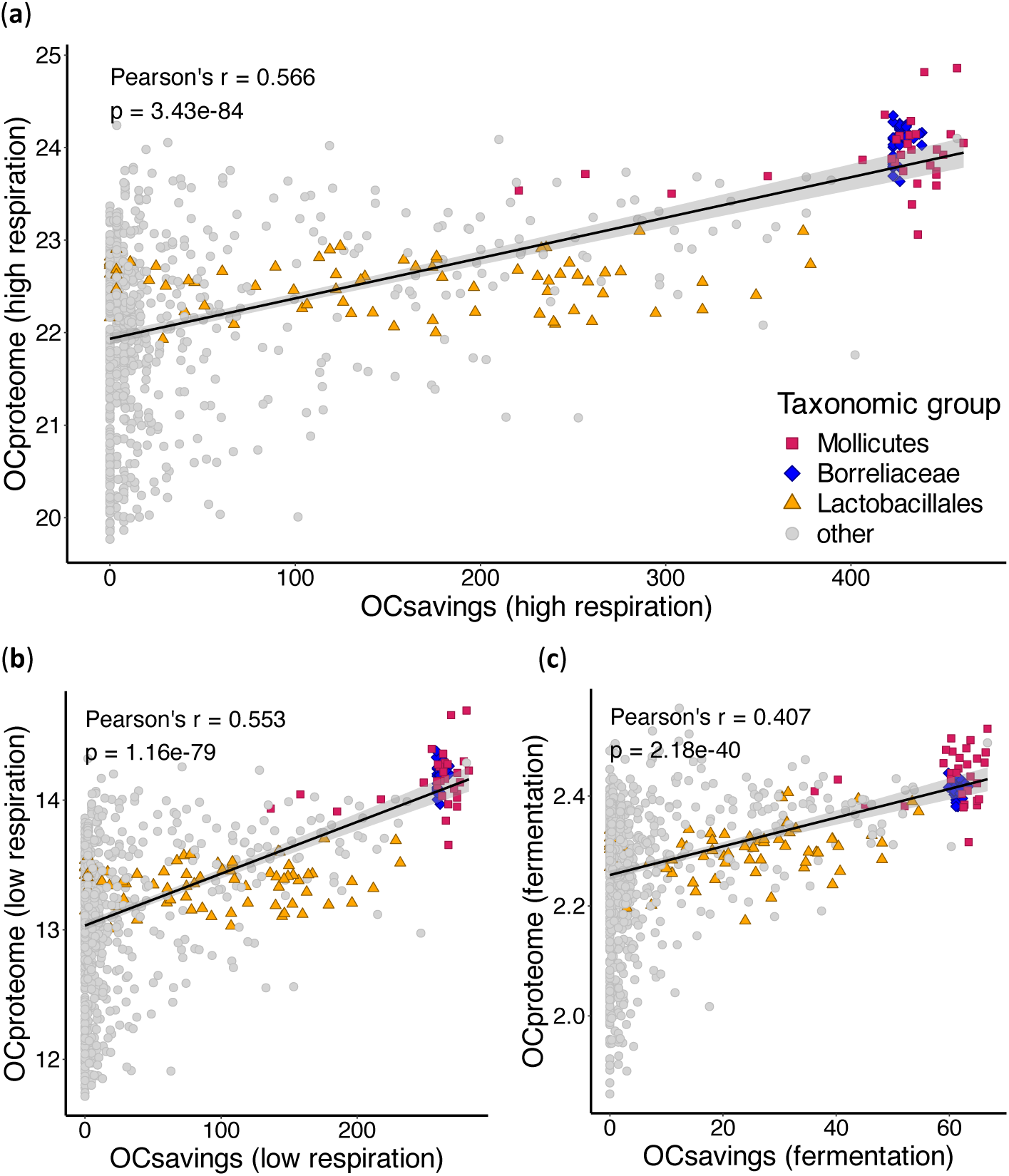
Correlation between the average AA cost per proteome and the amount of outsourced energy for AA biosynthesis. We estimated the completeness of AA biosynthesis pathways for 980 bacterial species. For each proteome, we calculated its opportunity cost (OC_*proteome*_) by multiplying the opportunity cost of each AA by its frequency in the proteome and summing the resulting values (see Materials and Methods). For each species, we also estimated the energy saved by outsourcing AA biosynthesis (OC_*savings*_). This was calculated by multiplying the auxotrophy index (AI) by the opportunity cost for each AA and summing the values across all 20 AAs (see Materials and Methods). Opportunity costs for each AA were previously estimated for three different respiratory modes: (**a**) high respiration, (**b**) low respiration, and (**c**) fermentation [2]. The Pearson correlation coefficient (r) and p-value are displayed on the graph.

### 2.4. Expensive proteins have ecologically relevant functions

To investigate the functional background of the most expensive proteins in Mollicutes, Borreliaceae, and Lactobacillales, we conducted an enrichment analysis of COG functions. Within each of these three clades, we separately calculated the opportunity cost of every protein (Equation 4) and used MMseqs2 clustering [17] to group them into homologous clusters [1,2]. Next, we determined the opportunity cost of each cluster by averaging the opportunity costs of its members (Equation 5). Finally, we performed an enrichment analysis of COG functions for the top 10%, 20%, 30%, 40%, and 50% most expensive clusters within each clade (Fig. 4).

**Figure 4.**
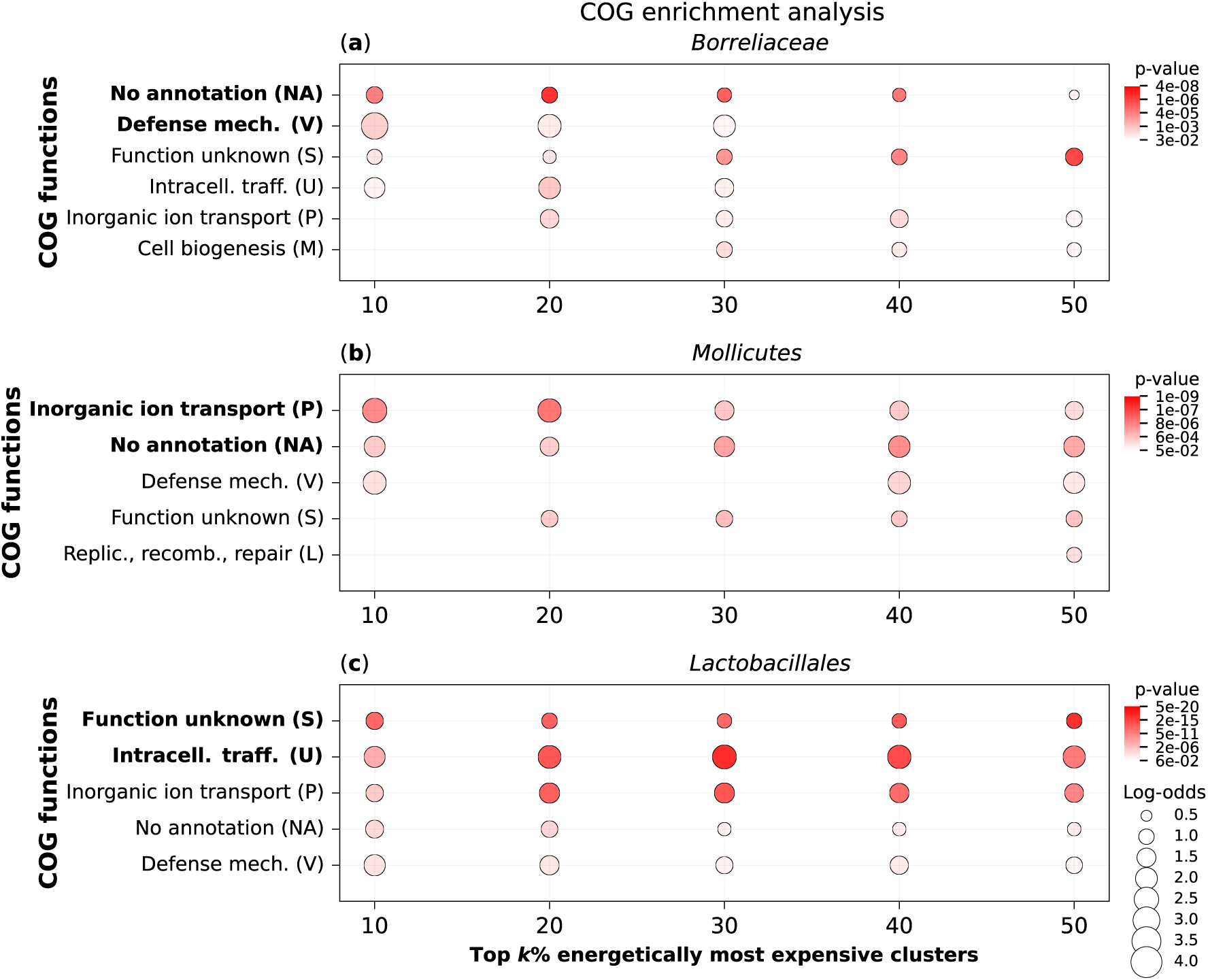
COG Enrichment analysis of the most expensive protein clusters in highly auxotrophic bacterial groups. We analyzed the proteomes of (**a**) Borreliaceae (50 species), (**b**) Mollicutes (31 species), and (**c**) Lactobacillales (74 species) from our full database of 980 bacterial species. Each dataset was clustered separately using the MMseqs2 algorithm to identify clusters of homologous proteins. COG functions were assigned to each cluster using EggNOG-mapper (see Materials and Methods). Clusters without functional annotations were labeled as NA (i.e., no annotation). We performed an overrepresentation analysis for the top k% of clusters ranked by OC_*cluster*_ (Equation 5), with k = 10, 20, 30, 40, and 50, using a one-tailed hypergeometric test. The resulting p-values were corrected for multiple testing using the Benjamini–Hochberg method. Only enrichment signals with p-values < 0.05 are shown (Table S4).

All three bacterial clades show enrichment in a similar set of COG functions; however, the statistical significance of these enrichments varies between the clades. For instance, proteins with unknown functions are among the top two functional categories with the most significant enrichments (i.e., the lowest p-values). However, the second most significantly enriched category differs across clades: defense mechanisms in Borreliaceae (Fig. 4a), inorganic ion transport in Mollicutes (Fig. 4b), and intracellular trafficking in Lactobacillales (Fig. 4c). This suggests that many of the most expensive proteins in these bacterial clades remain severely understudied, while those that have been characterized are involved in different aspects of cellular processes.

## 3. Discussion

In our previous work, we applied the concept of functional outsourcing [1] to amino acid (AA) biosynthesis in animals and developed a model describing the conditions required for the evolution of AA auxotrophies [2]. This model predicts that the loss of AA production capabilities is not random because energy-optimizing selection favors the loss of pathways for energetically costly AAs. As a result, AA auxotrophs are able to more freely explore protein sequence space by reducing selective constraints on the use of expensive AAs in their proteomes [2]. To test the broader applicability of this model across major clades, we investigated the patterns of AA auxotrophies in bacteria.

While animals exhibit nearly identical sets of auxotrophies among each other, bacterial metabolisms are far more diverse, making bacterial auxotrophies more challenging to detect and interpret [4,7,9]. Although it is known that bacterial AA auxotrophies can confer a fitness advantage in competition with prototrophic strains [7,10], it remains unclear which adaptive benefits are gained through AA auxotrophy and whether they are linked to energy management [2].

Our results reveal a significant positive correlation between the AA auxotrophy index averaged across tested bacteria and the AA opportunity cost, as predicted by the AA outsourcing model [2]. This finding aligns with earlier observations that costlier AAs promote stronger microbial cross-feeding interactions [11] and that AA auxotrophies confer a fitness advantage [7,10]. Beyond our study, only one prior investigation has explicitly examined bacterial AA auxotrophies from an energy-usage perspective, reporting no significant correlation between the frequency of an AA being auxotrophic and its biosynthetic cost [4]. The reasons for this discrepancy remain unclear and may stem from differences in bacterial genome datasets, auxotrophy detection pipelines, biosynthetic cost estimates, or uneven sampling of bacterial groups. Nonetheless, other trends observed in that study [4] closely resemble our findings— tryptophan, leucine, histidine, valine, and serine are commonly auxotrophic.

To further test the robustness of our results, we analyzed data from another comparable study that estimated AA auxotrophies in the gut microbiome using metabolic modeling, though it did not include energy calculations [14]. Similar to our study, this dataset also showed a significant correlation between the frequency of an AA being auxotrophic and its biosynthetic cost, with an even higher correlation coefficient than in our results (Figure S1). Consistent with our findings, they reported that tryptophan, the most expensive AA, is the most commonly auxotrophic. Moreover, their auxotrophy profiles closely resemble ours, despite being derived from a niche-derived community—gut microbiome bacteria [14]. Interestingly, they also found that host-microbiome and microbe-microbe interactions can play a crucial role in the maintenance and spread of AA auxotrophy [14], aligning with our concept of functional outsourcing [1]. Collectively, these findings suggest that, in at least some bacterial groups, AA auxotrophies are influenced by energy savings at the level of AA biosynthesis.

The second prediction of our model is that bacterial species with more auxotrophies should have more expensive proteomes. Our results confirm this prediction, showing that auxotrophic species indeed maintain genes that encode more expensive proteomes. This finding suggests that AA auxotrophy fundamentally influences protein evolution. The energy savings achieved through outsourcing AA biosynthesis relax the constraints on incorporating costly AAs into proteins, thereby enabling auxotrophic organisms to explore protein sequence space more freely [1,2,18–20].

This is best illustrated by Borreliaceae and Mollicutes, which exhibit extreme levels of both proteome expensiveness and auxotrophy. They have outsourced most of their metabolic processes to their hosts [21–23], making their AAs relatively inexpensive, which likely facilitated pathogen-host coevolution. In contrast, some Lactobacillales species exhibit only a few auxotrophies while others display many, yet this variation does not impact proteome costs, which remain relatively constant. This pattern most likely arises because Lactobacillales predominantly rely on fermentative metabolism [24]. As predicted by our model [2], fermentation lowers AA synthesis costs, thereby relaxing the selective pressures that drive the outsourcing of expensive AA production. This suggests that auxotrophies in Lactobacillales are more strongly determined by the availability of specific AAs in the environment rather than by the energy burden of AA biosynthesis [2].

Based on these findings, we hypothesize that the increase in the frequencies of costly AAs in auxotrophic species could result in the evolution of proteins with novel functions [1,2,18–20]. Interestingly, the most auxotrophic groups with the most expensive proteomes in our analysis are Borreliaceae and Mollicutes, the members of which are notorious for causing severe infections that are difficult to manage [25–27]. It is also known that *Borreliella* (syn. *Borrelia*) *burgdorferi* (Spirochaetales) harbors many Borreliaceae-specific genes with unknown functions, which may be implicated in the development of Lyme disease [28].

Our enrichment analysis further supports the idea that auxotrophies facilitate the evolution of proteins with novel functions. In all three tested groups, many of the most expensive protein clusters have unknown functions, suggesting that the energy saved through outsourcing is invested in lineage-specific proteins. In Borreliaceae, the enrichment of defense-related functions among expensive proteins may be associated with their complex immune evasion strategies [29,30]. Similarly, inorganic ion transport functions are the most expensive category in Mollicutes, whose pathogenicity is closely linked to alterations in the ion transport of host cells [31]. Finally, the enrichment of expensive intracellular trafficking functions in Lactobacillales likely reflects their ability to acquire nutrients through symbiotic interactions [24]. Taken together, our results suggest that auxotrophy-related energy shifts may drive the evolution and function of lineage-specific genes, a topic that warrants further in-depth investigation.

In conclusion, our results suggest a global macroevolutionary trend in bacteria where AA biosynthesis capability is shaped by energy-saving selection, ultimately leading to the evolution of more expensive proteomes. However, bacteria exhibit remarkable ecological and metabolic diversity, thriving in a variety of physically and chemically unique habitats [9,32,33]. This diversity raises the possibility that factors beyond energy costs influence reductions in AA biosynthetic pathways. Future studies focusing on specific bacterial lineages and ecologies will be crucial for uncovering the role of such factors in shaping the evolution of AA biosynthetic capabilities.

## 4. Materials and Methods

### 2.1. Databases, completeness score and auxotrophy index

To create the database of bacterial proteomes (Table S1), we combined datasets previously assembled for the phylostratigraphic analyses of *Bacillus subtilis* [34] and *Borrelia burgdorferi* [28], along with a resolved phylogeny of *Escherichia coli*. This resulted in a database of 980 bacterial proteomes, predicted from their corresponding genomes, representing most major bacterial lineages. The proteomes, primarily retrieved from the NCBI database and supplemented by the Ensembl database, were evaluated for contamination using BUSCO [35], and all were confirmed to be free of contamination [1].

In our previous study, we retrieved pathways and enzyme codes involved in AA biosynthesis from the KEGG and MetaCyc databases [2,36]. For AAs that can be synthesized via multiple alternative pathways, we treated each pathway separately, even when they shared some enzymes. Using this collection of enzyme codes associated with AA biosynthesis, we retrieved bacterial protein sequences from the KEGG database [37,38]. For each genus with representatives annotated in KEGG, we selected the species with the largest number of enzymes catalyzing reactions in AA biosynthesis pathways. This process resulted in a protein sequence reference database comprising 387,892 AA biosynthesis enzymes across 2,095 species (Table S2).

In the next step, we combined the downloaded enzyme sequences known to be involved in AA biosynthesis pathways with all sequences from our bacterial proteomes (4,230,625 sequences across 980 species) into a single database. We then clustered this combined database using MMseqs2 [17] with the following parameters: -cluster-mode 0, -cov-mode 0, -c 0.8, and –e 0.001. Clustering with these parameters generated clusters whose members exhibit highly similar architectures, as the alignment between query and target sequences covered at least 80% of their length [1].

Using the presence of enzymes involved in AA synthesis, we functionally annotated the remaining members of each cluster by transferring functional information from known AA synthesis enzymes. For each AA biosynthesis pathway and species in the database, we calculated a pathway completeness score (CS) by dividing the number of detected enzymes by the total number of enzymes in that pathway, resulting in values ranging from 0 to 1. If a species contained alternative biosynthetic pathways for an AA, the pathway with the highest completeness score was selected. We then calculated the AA auxotrophy index (*AI*_*i*_) using the following equation:

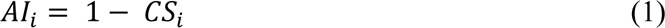

In this equation CS is completeness score and *i* denotes one of 20 AAs (*i =* 1, …, *20*).

To evaluate the performance of our method, we utilized a previously established testing set of experimentally identified prototrophies and auxotrophies [4,14] to estimate pathway completeness scores (CS). The first tested dataset comprised 160 fully prototrophic species [4]. Our approach exhibited an error rate of 0.012, indicating that approximately 1.2% of amino acids were incorrectly identified as auxotrophic (Data S1). The second dataset included 15 species with at least one known auxotrophy [14]. In this case, we detected an error rate of 0.188 for false prototrophs, meaning that around 19% of amino acids were incorrectly classified as prototrophic (Data S2). Taken together, these error rates suggest that our approach to auxotrophy detection is conservative and aligns with error rates reported for similar methods [4,14].

### 2.2. Opportunity cost measures

In our earlier study, we calculated the opportunity cost of biosynthesis for each AA depending on the three respiration modes: high respiration, low respiration and fermentation [2]. The opportunity cost is calculated as the sum of the energy lost in the synthesis of AAs and the energy that would have been produced if a cell catabolized precursors instead of making AAs [2,39,40]. This measure reflects the overall impact of AA synthesis on the cell’s energy budget, and is rather consistent regardless of the carbon source used by the bacteria [41]. To approximate how much ATP is generated from the reducing equivalents in different respiratory conditions, we converted the reducing equivalents to ATP in the following way: (i) ‘high respiration,’ representing fully functional oxidative phosphorylation: 1 NAD(P)H = 2 FADH2 = 2 ATP [12]; (ii) ‘low respiration,’ representing oxidative phosphorylation without proton pumping at complex I: 1 NAD(P)H = 2 FADH2 = 1 ATP, which corresponds, for instance, to the metabolism of *S. cerevisiae* and some *E. coli* strains [42,43]; (iii) ‘fermentation,’ representing anaerobic metabolism without conversion of reducing equivalents to ATP [2]. All of these values for each AA are available in Table S3.

Using the AA opportunity cost, we also calculated the opportunity cost of each proteome (*OC*_*proteome*_) using the following equation:

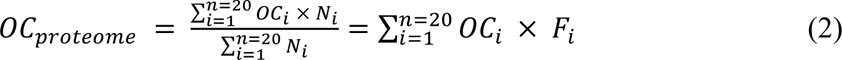

In this equation, *OC*_*proteome*_ is a weighted mean where *OC*_*i*_ represents the opportunity cost of the *i*-th AA, *N*_*i*_ denotes the total number of occurrences of this AA in the entire proteome, and *F*_*i*_ represents the frequency of the AA in the proteome (calculated as the number of occurrences of the AA divided by the total number of AAs in the proteome).

We also introduced a new measure (*OC*_*savings*_) to quantify the energy savings of a species by linking the AA auxotrophy index (*AI*_*i*_) to the opportunity cost. This measure estimates the energy saved by outsourcing AA production to the environment. It is calculated as follows:

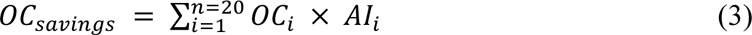

In this equation, *OC*_*i*_ denotes the opportunity cost of a given AA, while *AI*_*i*_ denotes the AA auxotrophy index.

### 2.2. COG functions enrichment analyses

For functional analyses, we analyzed separately the proteomes of Borreliaceae (50), Mollicutes (31), and Lactobacillales (74) taken from our full database of 980 bacterial species. We clustered the three datasets separately using the MMseqs2 cluster algorithm (the 14-7e284 version) with the following parameters: -e 0.001 -c 0.8 --max-seqs 400 --cluster-mode 1 [1,17]. For each protein in the datasets we obtained its COG annotations using the EggNOG-mapper (version 2.1.12) [44] with the diamond (version 2.1.8) searching tool [45]. We also computed for each protein its opportunity cost (*OC*_*protein*_) for high respiration mode [2] as:

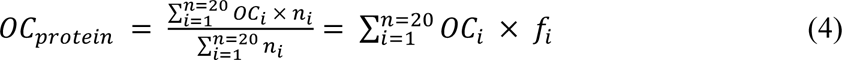

In this equation, *OC*_*protein*_ is a weighted mean where *OC*_*i*_ denotes the opportunity cost of the *i*-th amino acid, *n*_*i*_ is the number of occurrences of the *i*-th of amino acid in a protein and *f*_*i*_ is the frequency of the *i*-th amino acid in a protein.

Finally, we performed the functional enrichment analysis of the three datasets independently. For each dataset, a cluster was assigned a COG function if at least one of its members was annotated with that function. The clusters whose proteins had no annotations were assigned with NA. The enrichment analysis was done for clusters with at least 10 members. For each cluster we calculated the average opportunity cost of a cluster (*OC*_*cluster*_) as follows:

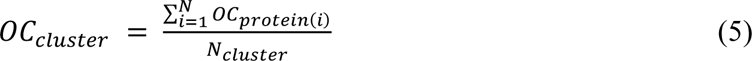

In this equation, *OC*_*protein(i)*_ is the opportunity cost of the *i*-th protein in a cluster (see Equation 4) and *N*_*cluster*_ is the number of proteins in the cluster.

We performed an overrepresentation analysis for the top k% of clusters ranked by OC*cluster*, with *k* = 10, 20, 30, 40, 50, using a one-tailed hypergeometric test as implemented in the Python *scipy.stats* module. The obtained p-values were corrected for multiple testing and adjusted using the Benjamini–Hochberg method as implemented in the Python *statsmodels* library [46]. All results of enrichment analysis are shown in Table S4.

To calculate correlations, we used the cor.test() function in the R stats (v. 3.6.2) package. The heatmap was visualized using the ggtree R package [47].

## Supplementary Materials

The following supporting information can be downloaded at: www.mdpi.com/xxx/s1, Table S1: Bacterial database; Table S2: Enzyme reference database; Table S3: Amino acid biosynthesis costs; Table S4: Results of enrichment analysis; Data S1: Prototrophy evaluation; Data S2: Auxotrophy evaluation; File S1: Fully resolved tree with AA auxotrophy heatmap; Figure S1: Correlation between AA biosynthesis cost and the average AA auxotrophy index for Starke et al. 2023 data.

## Author Contributions

Conceptualization, N.K., T.D.-L. and M.D.-L.; methodology, N.K., T.D.-L. and M.D.-L.; software, N.K. and M.D.-L.; validation, N.K., T.D.-L. and M.D.-L.; formal analysis, N.K., T.D.-L. and M.D.-L.; writing—original draft preparation, N.K., T.D.-L. and M.D.-L.; writing—review and editing, N.K., T.D.-L. and M.D.-L.; visualization, N.K.; supervision, T.D.-L. and M.D.-L. All authors have read and agreed to the published version of the manuscript.

## Data Availability Statement

All data are available in the supplementary materials.

## Acknowledgments

We thank M. Futo, A. Tušar, S. Koska, D. Franjević and G. Klobučar for discussions. We used the computational resources of the University Computing Center - SRCE (Padobran) and the Institute Ruđer Bošković.

---

**Figure S1.**
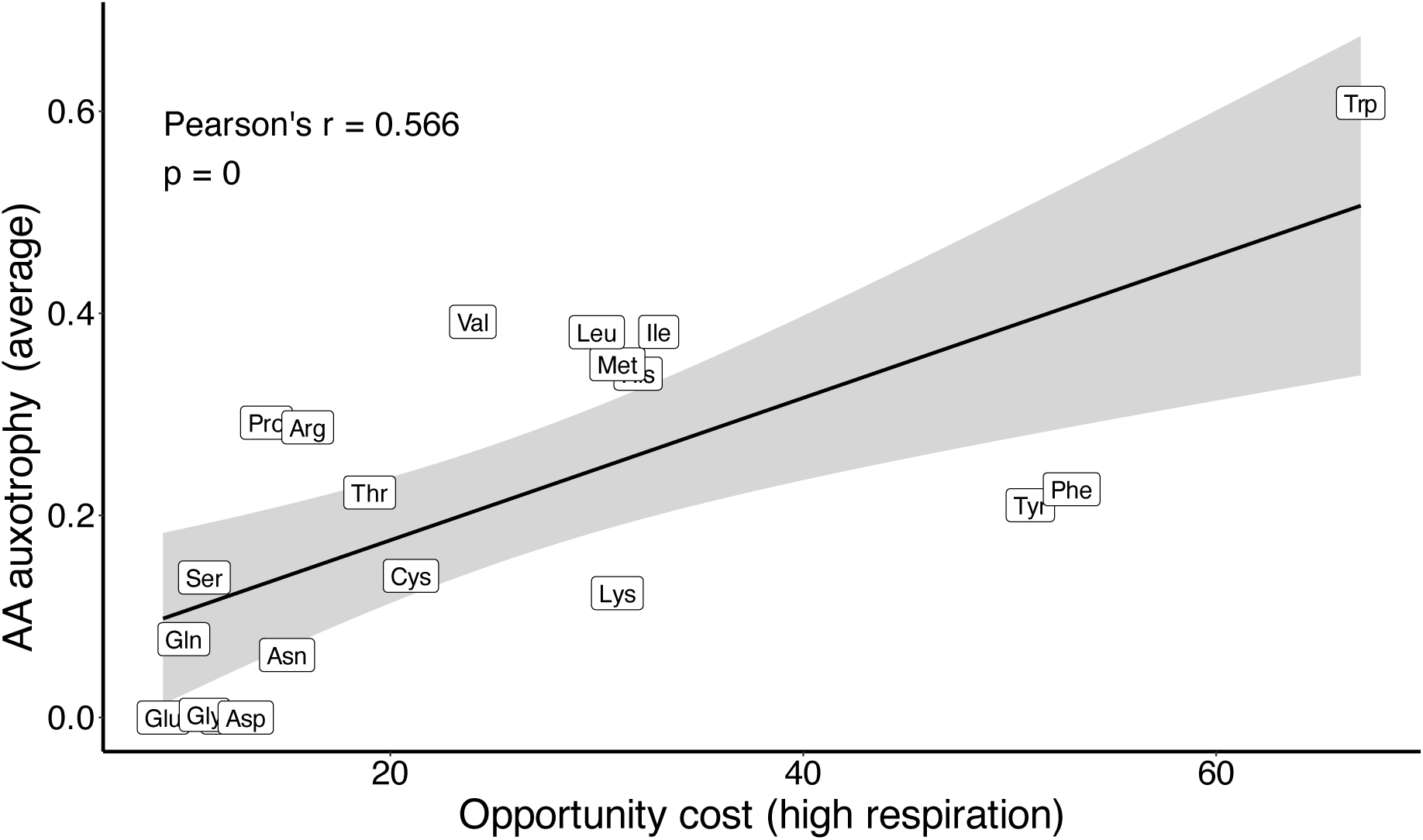
Correlation between AA biosynthesis cost and the average AA auxotrophy based on data from Starke et al. (2023). We calculated the average of AA auxotrophy measures provided in the Starke et al. (2023) study for 3687 bacterial species. We correlated this value with the opportunity cost of each AA, calculated for high respiration mode (see Materials and Methods). Pearson correlation coefficient and p-value are shown on the graph.

